# Selfish migrants: How a meiotic driver is selected to increase dispersal

**DOI:** 10.1101/2021.02.01.429134

**Authors:** Jan-Niklas Runge, Hanna Kokko, Anna K. Lindholm

## Abstract

Meiotic drivers are selfish genetic elements that manipulate meiosis to increase their transmission to the next generation to the detriment of the rest of the genome. One example is the *t* haplotype in house mice, which is a naturally occurring meiotic driver with deleterious traits—poor fitness in polyandrous matings and homozygote inviability or infertility—that prevent its fixation. Recently, we discovered and validated a novel effect of *t* in a long-term field study on free-living wild house mice and with experiments: *t*-carriers are more likely to disperse. Here, we ask what known traits of the *t* haplotype can select for a difference in dispersal between *t*-carriers and wildtype mice. To that end, we built individual-based models with dispersal loci on the *t* and the homologous wildtype chromosomes. We also allow for density-dependent expression of these loci. The *t* haplotype consistently evolves to increase the dispersal propensity of its carriers, particularly at high densities. By examining variants of the model that modify different costs caused by *t*, we show that the increase in dispersal is driven by the deleterious traits of *t*, disadvantage in polyandrous matings and lethal homozygosity or male sterility. Finally, we show that an increase in driver-carrier dispersal can evolve across a range of values in driver strength and disadvantages.

## Introduction

Conflict is everywhere. It not only takes place between species or between individuals, but also within individuals (Burt and Trivers 2006; Queller and Strassmann 2018). In general, all genetic elements are selected to increase the frequency at which they are copied to future generations. Most elements achieve this by increasing the fitness of the organism that carries them, which aligns the interests of the organism and its genome. However, there are also elements that increase their own representation in future generations at the expense of the rest of the genome, without a positive fitness effect on the organism. Since some even cause harm to the organism’s fitness, these elements are known as selfish genetic elements (Burt and Trivers 2006). Selfish genetic elements come in a variety of forms. For example, killer meiotic drivers increase their frequency in the functioning gametes of an organism by inhibiting or destroying gametes that do not carry the driver (Núñez et al. 2018).

Killer meiotic drivers work in one of two ways (Núñez et al. 2018): Either they release a killer element that attacks a target locus in *trans* (on the homologous chromosome), or they create a poison that attacks all meiotic products (indiscriminate of whether they carry the driver), together with an antidote that acts in *cis* and thus rescues only driver-carrying meiotic products. These poison-antidote drivers commonly work only in males and could cause reduced sperm competitiveness for reasons such as imperfect rescue and reduced gamete counts (Price and Wedell 2008). In such a scenario, the fitness outcomes for the driver differ dramatically between monandrous and polyandrous matings. In the latter case, where sperm from multiple males compete over fertilization, the driver-carrying poor sperm competitors are at a significant disadvantage.

Females commonly mate with multiple males in wild populations (Taylor et al. 2014). The frequency of polyandry has been linked to genetic and environmental factors, e.g. male fertility (Sutter et al. 2019), local density (Dean et al. 2006; Firman and Simmons 2008; Manser et al. 2020), and presence of meiotic drivers (Price et al. 2008). Less is known about meiotic drivers themselves adapting to local variation in polyandry: if drivers do better in single matings, can they somehow avoid ending up in polyandrous situations? One possibility is that drivers could increase the dispersal propensity of their carriers. This hypothesis is based on the argument that dispersal via movement to less dense populations on average may bring driver-carriers to areas with less polyandry. Dispersal could also help avoid matings with another driver carrier; such matings cause some offspring to be homozygous for the driver, which is detrimental in several drive systems (Lewontin and Dunn 1960; Larracuente and Presgraves 2012; Fishman and Kelly 2015).

We investigate these possibilities for a naturally occurring poison-antidote male meiotic driver in house mice (*Mus musculus*), the *t* haplotype, for which there is a wealth of knowledge of its traits, but we also extend our work to varying key traits in order to generalize the conclusions. The *t* haplotype comprises a 35 Mb linked region on an autosome, estimated to be two million years old (Silver 1993; Kelemen and Vicoso 2018). It manipulates spermatogenesis to increase its own chances of transmission (Lindholm et al. 2019; Charron et al. 2019; Amaral and Herrmann 2021). Heterozygous (notation: +/*t*) males transmit the *t* haplotype with 90% probability, leaving only 10% for the homologous wildtype chromosome (denoted +). This marked contrast with the ‘fair’ Mendelian rate of transmission of 50% makes the *t* “selfish”.

Despite enjoying a transmission advantage, *t* does not fix or persist at high frequencies in natural populations (Ardlie and Silver 1998). One reason is that homozygous (*t*/*t*) carriers of the *t* haplotype are either inviable (Klein et al. 1984) or sterile as males (Lyon 1986), which is a large cost to the *t*’s fitness (Dunn and Levene 1961; Safronova 2009; Lindholm et al. 2013; Sutter and Lindholm 2015). The *t* is however even less frequent in natural populations than would be predicted based on its homozygous disadvantages (Bruck 1957; Ardlie and Silver 1998), a pattern known as the “*t* paradox” (Manser et al. 2011). This paradox was explained by another deleterious trait of the *t*: the sperm of *t*-carrying males (+/*t*), while almost exclusively transmitting the *t*, are less competitive than sperm of wildtype (+/+) males (Sutter and Lindholm 2015; Manser et al. 2017). Consequently, +/*t* males sire a clear minority (11-19%) of the offspring of polyandrous matings when in competition with +/+ males.

An increased dispersal propensity conceivably improves the *t*’s chances of being present in multiple populations, new populations, and populations in which it is (temporarily) fitter than the wildtype (Levin et al. 1969; Hamilton and May 1977; Comins et al. 1980). In general, dispersal leaves more resources for related kin (Hamilton and May 1977) (in this case, other *t* alleles). This might not promote dispersal of *t* above that of the wildtype *per se*, since +/+ enjoy equivalent benefits as well (likewise, arguments such as ‘being present in multiple populations is beneficial’ apply to +/+ too), but for *t* there is a unique benefit of leaving a *t*-rich habitat patch. Their departure counteracts the possibility of two philopatric +/*t* individuals mating with each other and producing inviable or infertile *t*/*t* offspring. If dispersal of *t* brings its carrier to a population with a lower *t* frequency, the benefit occurs both at the new as well as the natal site.

As a flipside, however, entering dense, +-rich habitat patches induces a larger risk of losing out in sperm competition, because the frequency of polyandrous matings increases with population density in house mice (Dean et al. 2006; Firman and Simmons 2008; Manser et al. 2020). This makes us hypothesize that net selection on *t*-associated density-dependent dispersal will depend on polyandry. On the other hand, if the homozygous costs are larger than the polyandrous disadvantage, then we would expect +/*t* to disperse preferentially out of low density sites (where *t* frequency is expected to be high due to the effectiveness of the meiotic drive).

In this framework, *t* behaves somewhat like an infection (Lion et al. 2006), though for unique reasons (homozygote inviability). If *t* increases in frequency as a result of successfully entering new local populations, the relative fitness of *t* will decrease over time. Whether this selects for dispersal even out of low density habitats (with low multiple mating frequencies), depends on the balance of all the costs and benefits of dispersal. As a whole, we expect the costs and benefits of dispersal to differ between the wildtype and the *t* haplotype. Indeed, our previous empirical work on free-living wild house mice found that *t*-carrying juveniles were more likely to emigrate, and were over-represented in migration events (Runge and Lindholm (2018), see Figure 1) as well as among dispersers in experimental setups (Runge and Lindholm 2021).

**Figure 1:**
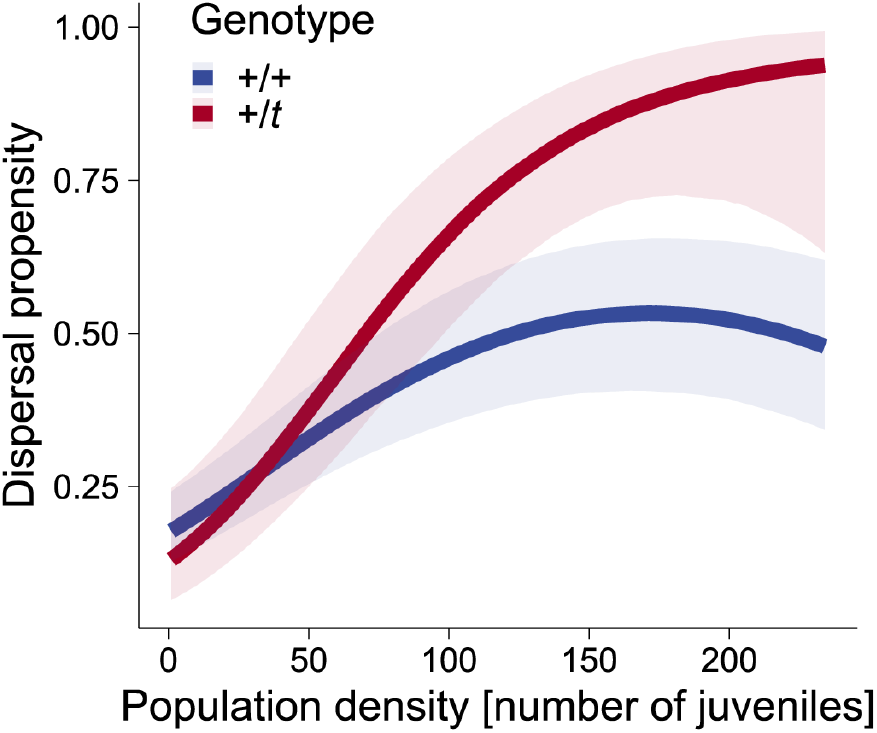
Differences in juvenile dispersal propensity between +/*t* and +/+ in a long-term field study, replotted from Runge & Lindholm (2018).

In this study we present results from individual-based models that simulate the evolution of the *t* haplotype’s influence on its carrier’s dispersal propensity. The results provide quantitative support for the hypothesis that *t* should evolve a density-dependent increased dispersal propensity. By considering multiple hypothetical scenarios, we find the *t*’s disadvantages in homozygous viability and sperm competition to be the main drivers of *t*’s elevated and density-dependent dispersal propensity. Additionally, we show that the result requires moderate spatial heterogeneity in density. Similar results arose in simulated *t* variants that produce viable but infertile *t*/*t* males, and more generally, we found hypothetical drivers with weak drive and low fitness costs to increase dispersal across a broad range of trait values.

## The model

The simulations were written and executed in NetLogo 6.1.1 (Wilensky 1999) and analyzed with R 3.6.1 with packages data.table 1.12.2 (Dowle and Srinivasan 2019), dplyr 0.8.5 (Wickham et al. 2019), ggplot2 3.1.1 (Wickham 2009), readr 1.3.1 (Wickham et al. 2018), and stringr 1.4.0 (Wickham 2019).

**Table 1:**
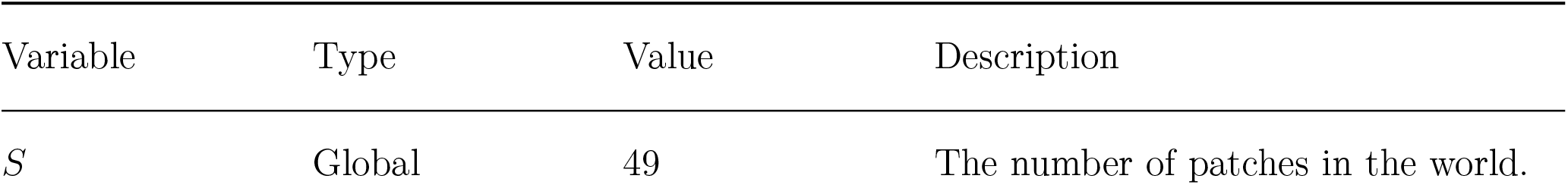
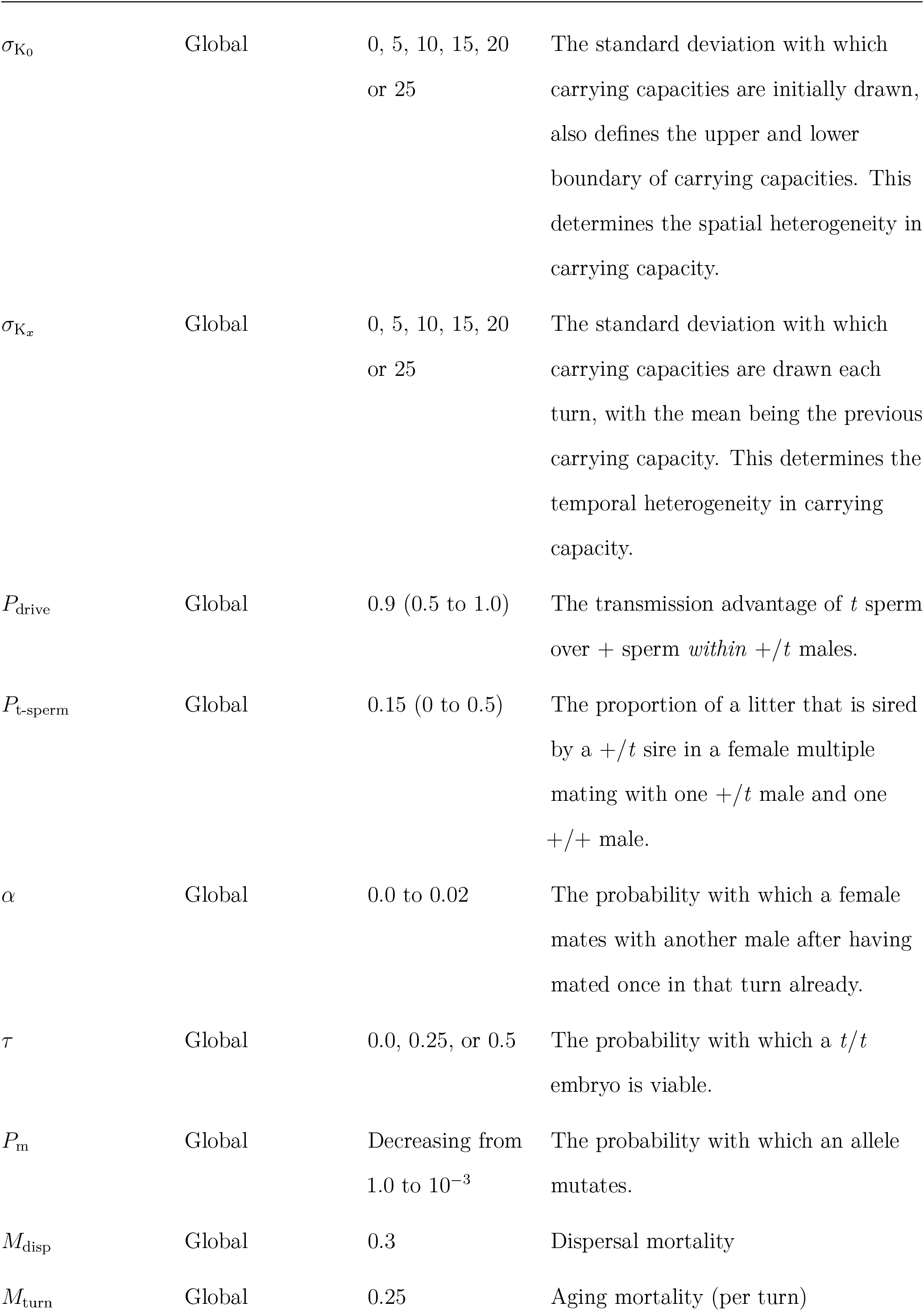
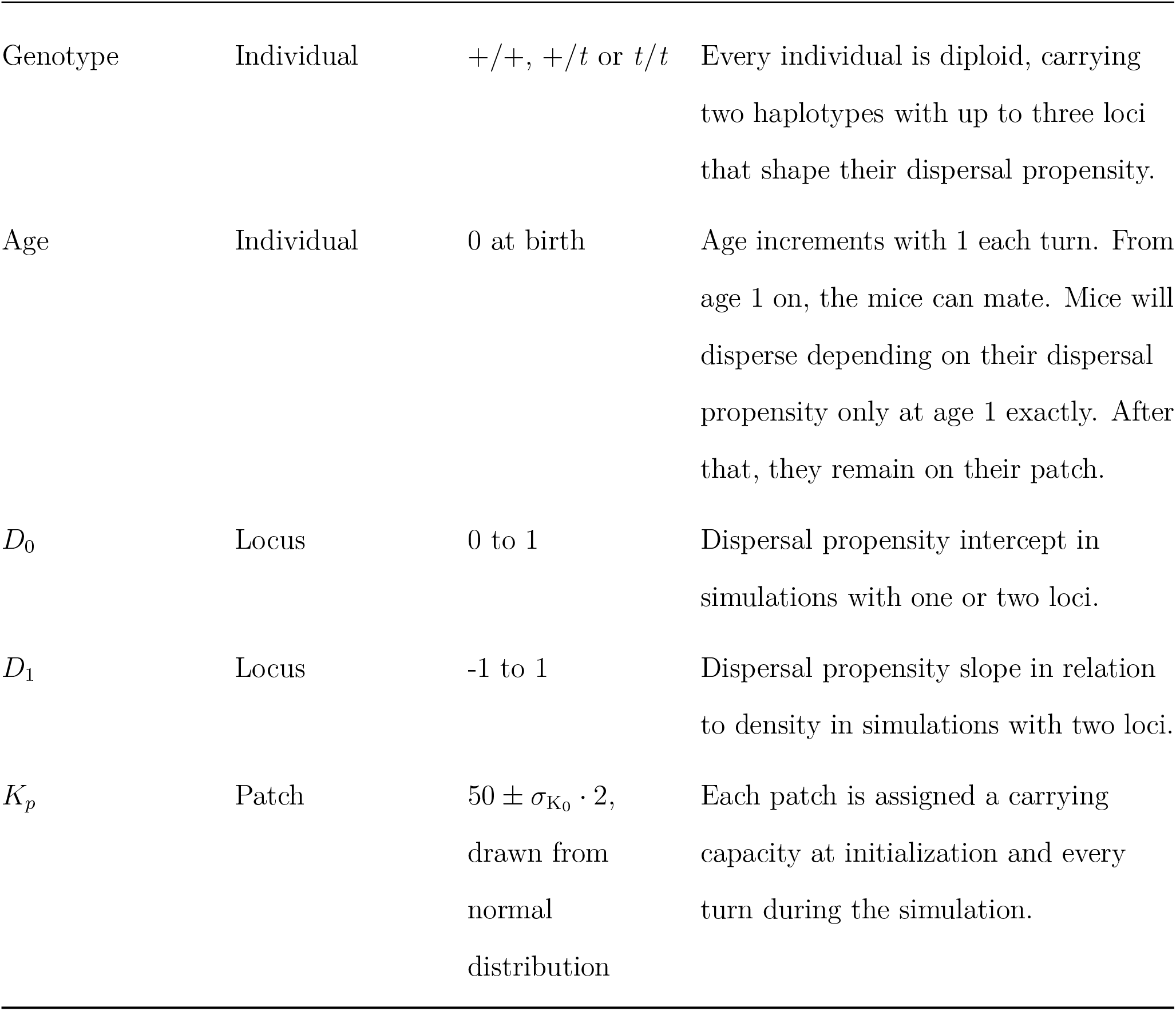
Overview of simulation variables

| Variable | Type | Value | Description |
| --- | --- | --- | --- |
| $S$ | Global | 49 | The number of patches in the world. |
| $\sigma_{K_0}$ | Global | 0, 5, 10, 15, 20<br>or 25 | The standard deviation with which carrying capacities are initially drawn, also defines the upper and lower boundary of carrying capacities. This determines the spatial heterogeneity in carrying capacity. |
| $\sigma_{K_x}$ | Global | 0, 5, 10, 15, 20<br>or 25 | The standard deviation with which carrying capacities are drawn each turn, with the mean being the previous carrying capacity. This determines the temporal heterogeneity in carrying capacity. |
| $P_{\text{drive}}$ | Global | 0.9 (0.5 to 1.0) | The transmission advantage of $t$ sperm over $+$ sperm *within* $+/t$ males. |
| $P_{\text{t-sperm}}$ | Global | 0.15 (0 to 0.5) | The proportion of a litter that is sired by a $+/t$ sire in a female multiple mating with one $+/t$ male and one $+/+$ male. |
| $\alpha$ | Global | 0.0 to 0.02 | The probability with which a female mates with another male after having mated once in that turn already. |
| $\tau$ | Global | 0.0, 0.25, or 0.5 | The probability with which a $t/t$ embryo is viable. |
| $P_{\text{m}}$ | Global | Decreasing from<br>1.0 to $10^{-3}$ | The probability with which an allele mutates. |
| $M_{\text{disp}}$ | Global | 0.3 | Dispersal mortality |
| $M_{\text{turn}}$ | Global | 0.25 | Aging mortality (per turn) |
| Genotype | Individual | $+/+$, $+/t$ or $t/t$ | Every individual is diploid, carrying two haplotypes with up to three loci that shape their dispersal propensity. |
| Age | Individual | 0 at birth | Age increments with 1 each turn. From age 1 on, the mice can mate. Mice will disperse depending on their dispersal propensity only at age 1 exactly. After that, they remain on their patch. |
| $D_0$ | Locus | 0 to 1 | Dispersal propensity intercept in simulations with one or two loci. |
| $D_1$ | Locus | -1 to 1 | Dispersal propensity slope in relation to density in simulations with two loci. |
| $K_p$ | Patch | $50 \pm \sigma_{K_0} \cdot 2$,<br>drawn from<br>normal<br>distribution | Each patch is assigned a carrying capacity at initialization and every turn during the simulation. |

### World

The simulated world consists of *S* = 49 patches. Each patch *p* can carry *K*_*p*_ mice, with the carrying capacity *K*_*p*_ fluctuating over space and over time. The initial values for *K*_*p*_ are drawn from a normal distribution with a mean of 50 and a standard deviation of 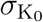. Each turn (generation), *K*_*p*_ is then re-drawn from a normal distribution with a mean of the patch’s current carrying capacity and a standard deviation of 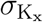; this procedure creates temporal autocorrelation between future and current densities. However, carrying capacities are re-drawn until they are within 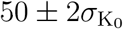 to prevent unbounded drift towards extreme values. These two *σ* represent the spatial 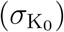 and temporal 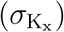 heterogeneity of the environment.

Dispersal is the act of changing the patch and there is no other way to move between the patches, effectively making them islands. Mice that share a patch have indistinguishable physical locations.

### The population

The model tracks the haplotypes of diploid individuals of differing sex and age (overview: Table 1). An individual carries two homologous chromosomes. Each chromosome is a haplotype that comprises two or three linked loci, which differs based on whether we simulated density-dependent or density-independent dispersal propensities. One locus determines whether the haplotype is + or *t*. Thus, an individual can be +/+ or +/*t*, with *t*/*t* not fully viable or fertile in some simulation conditions. The other loci determine the dispersal phenotype as described in *Dispersal* below. An individual’s age is the number of turns (see below) since birth.

### Initialization

At the beginning of each simulation, 5,000 mice of age 1 are placed randomly onto patches. Initially, all mice are +/+ and 50% are female. All mice that are placed onto patches start with a propensity to disperse between 0 and 1 (uniformly drawn), independent of density (i.e. 0 *≤ D*_0_ *≤* 1; *D*_1_ = 0, see below). After 10% of the simulation’s turns, 50% of all mice in the simulation, chosen randomly, are converted from +/+ to +/*t* by converting one chromosome from + to *t* while keeping all dispersal loci values of that chromosome unchanged. The temporal delay in placing the *t* haplotype into the world is to ensure that the population has time to equilibrate to carrying capacity across the landscape before the competition between the + and the *t* allele begins, and to have proceeded past transient effects that are due to initializing the population with a wide range of dispersal propensities.

### Turns

Within each turn, the following procedures are run for all individuals sequentially (i.e. every procedure is done for all individuals before the next procedure begins): dispersal, mating, birth, and death together with aging of the survivors. Within each behavior, the order of individuals performing it is randomized.

### Behaviors

#### Dispersal

One or two loci, depending on the simulated scenario, determine the dispersal phenotype. In the one-locus models, only a density-independent propensity to disperse (*D*_0_) can evolve. In the two-locus models, *D*_0_ is the intercept and *D*_1_ is the slope of dispersal propensity in relation to density.

The intercept *D*_0_ can take values between 0 and 1 (0 and 100 percent (or percentage points) dispersal propensity), while the slope *D*_1_ can go from −1 to 1. Individuals evaluate the density of their patch at the beginning of a turn, and do not update it as some individuals begin to disperse; this ensures all individuals have the same information on the patch’s density.

One of the two haplotypes carried by each individual acts as dominant and only the values of the dominant alleles are used to determine the dispersal phenotype. In +/*t, t* is always dominant, while in +/+ and *t*/*t* one haplotype is chosen at random to be dominant.

A mouse’s dispersal propensity (at the patch’s current density) is evaluated against a uniformly drawn real number between 0 and 1 to determine whether the individual disperses. A dispersing focal mouse will move to a randomly chosen new patch (dispersal is global). Dispersal is also costly, leading to death with probability *M*_disp_.

#### Mating

In the mating procedure, focal females with age *≥* 1 are approached sequentially by all males of age *≥* 1 on the same patch, with the list of females ordered randomly every turn and the list of males ordered randomly for every female. The female will always mate with the first approaching male and mate with subsequent approachers with a probability of *α*, which creates a density-dependent increase in polyandry (SI Figure 1). Note that males are not limited in their mating (other than by females not always accepting them).

#### Birth

Each mated female begins her pregnancy with up to six offspring. The number of offspring *N*_*o*_ at the start of the pregnancy is affected by the local density in relation to the carrying capacity of the patch (reflecting increasingly constrained resources):

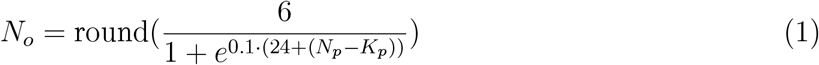

where *N*_*p*_ is the number of mice on the patch with carrying capacity *K*_*p*_. This relationship predicts zero offspring if the local population has reached carrying capacity, exactly one offspring at *N*_*p*_ = *K*_*p*_ *−* 1, and increasing numbers of offspring (up to 6) when densities fall clearly below the carrying capacity (see SI Figure 2). This assumption avoids flooding the population with unrealistically large numbers of offspring that would make up almost all of the population in the next turn after mice died due to density (see *Mortality* below).

**Figure 2:**
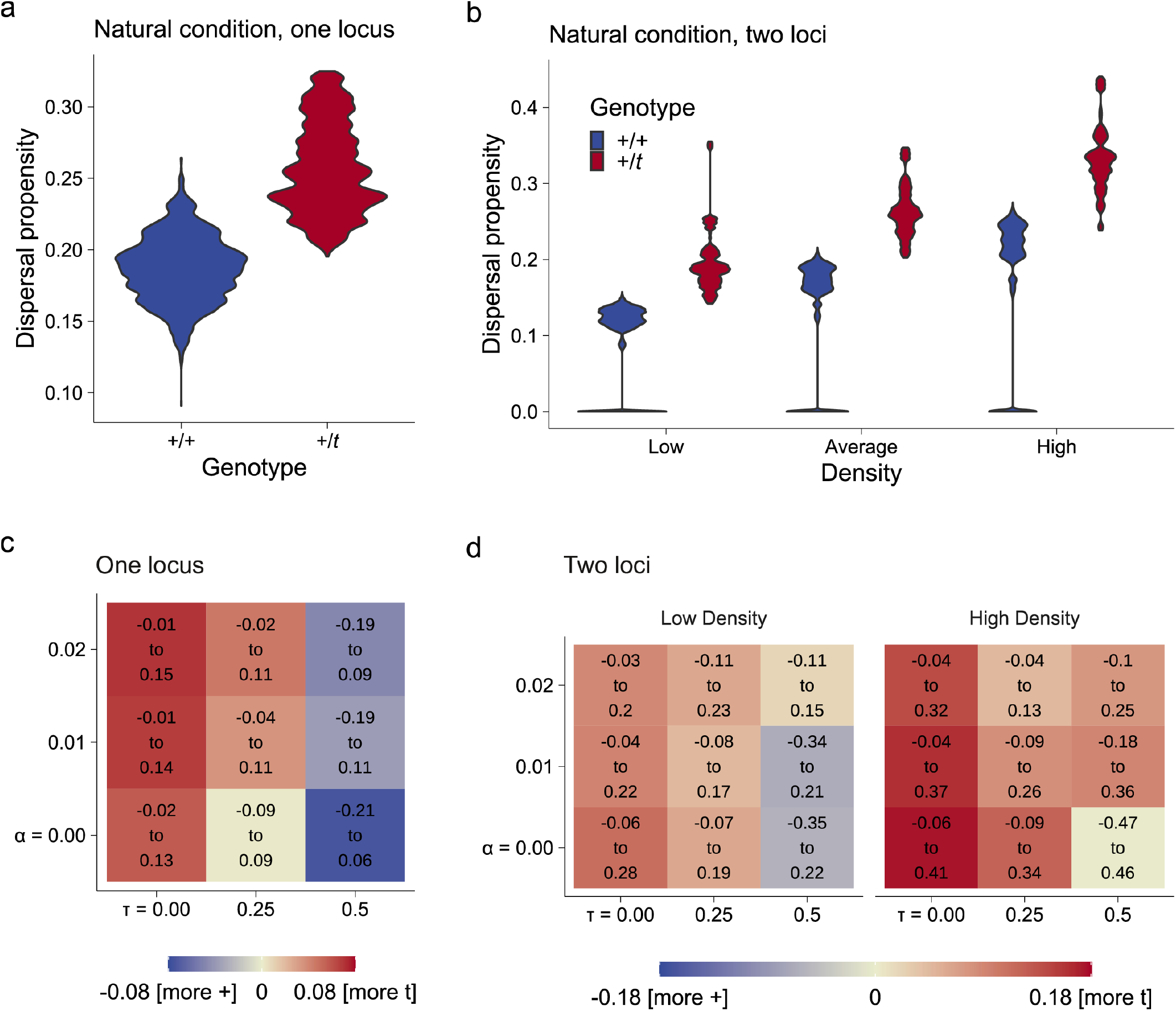
Overview of the differences in evolved dispersal propensities between +/+ and +/*t*. **a)** Violin plot of the evolved dispersal propensities of +/+ and +/*t* in the natural condition with one locus. **b)** Violin plot of the evolved dispersal propensities in two-locus models at different local densities. **c)** Heatmap showing the mean difference between *t* and + dispersal propensities in one locus models in varying polyandry *α* and *t*/*t* viability *τ*. Red indicates increased *t* dispersal, blue indicates increased + dispersal. The text indicates the 95% confidence interval. The natural condition of **a** is in the top left corner. **d)** Heatmap showing the mean difference between *t* and + dispersal propensities in two-locus models in low and high densities. The natural condition of **b** is in the top left corner.

Each offspring is assigned sex independently from each other (1:1 primary sex ratio); only viable offspring will be born, at which point they are assigned an age of 0. Embryonic viability is only affected in *t*/*t* homozygotes. A *t*/*t* will be viable with the probability *τ*, set globally for each simulation. The sire for each young is determined independently with the following procedure. For mothers who mated singly, the sire is obvious. For mothers who mated with multiple sires, every *t*-carrying mate has the probability of being a sire

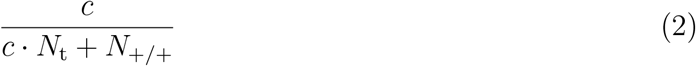

with the corresponding probability of a +/+ mate being the sire

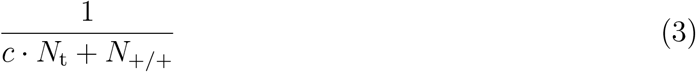

nlwhere *N*_t_ is the number of *t*-carrying (+/*t* or *t*/*t*) males that mated with that female, *N*_+/+_ is the corresponding number of +/+ males, and *c* represents the sperm-competitive ability of *t*-carrying males relative to +/+ males. To simplify interpretation of the results, we translated *c* into the probability of a *t*-carrying male being the sire when competing against one +/+ male *P*_t-sperm_, which is 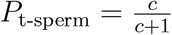, and we will refer to *P*_t-sperm_ from here. We chose *P*_t-sperm_ = 0.15 as the default, following experiments (Sutter and Lindholm 2015; Manser et al. 2017) that suggest values between 0.11 and 0.19.

If the sire is +/*t*, the *t*-carrying chromosome is transmitted with probability *P*_drive_ (and the +-carrying chromosome with the complementary probability 1 *− P*_drive_). Females, as well as +/+ and *t*/*t* males transmit a randomly chosen chromosome. We do not make the chromosomes recombine, thus all loci carried by a chosen chromosome are transmitted to the next generation.

Finally, the dispersal loci *D*_*i*_ mutate in the offspring with a probability that is initially higher (to allow for efficient searching of the space of possible reaction norms) and gradually diminishes. All dispersal loci mutate independently with a probability of 1.0 (100%) in the beginning, which linearly decreases to 10^*−*3^ over the first 10% of the turns, when only +/+ are in the simulation, and then goes back to 1.0 as *t* enters and decreases again to 10^*−*3^ over the final 90% of the turns. This temporal variation in mutation frequency is done to allow +/+ to find their optimum before *t* chromosomes enter and then give both + and *t* enough time and mutations to find an optimum for both. We found this approach to be better suited than an unchanging mutation rate, as genetic drift needs to be carefully balanced with mutations in our question due to the different effective population sizes of + and *t*.

In case a mutation takes place, the new value of the variable will be an addition or subtraction (chosen randomly) of 10^*−*3^ in case of the intercept *D*_0_ or of 2 *·* 10^*−*5^ in case of the slope *D*_1_. These values differ because of their different impact: a mutation of the slope increases or decreases the dispersal propensity at a density of 50 (mean carrying capacity) by 10^*−*3^, just as a mutation of the intercept does at a density of 0. Mutations that move the value outside the parameter space are set to the relevant boundary.

### Mortality

We include both density-independent and density-dependent mortality. A focal mouse dies due to density-independent causes with a probability of *M*_turn_ per turn. After applying this mortality to all mice, we further impose patch-specific carrying capacities *K*_*p*_ on the survivors, causing density-dependent mortality. The number of mice that die is *N*_*p*_ *− K*_*p*_, i.e. mice in excess of the carrying capacity are removed. The necessary mortality to achieve this is random with respect to traits or sex of the mice.

### Conditions

We refer to the set of values {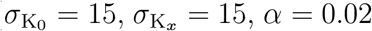, *α* = 0.02, and *τ* = 0} as the “natural condition” as it combines moderate temporal and spatial carrying capacity heterogeneity, frequent female multiple mating (*α* = 0.02) and fully lethal *t*/*t* (*τ* = 0), and leads to the evolution of realistic +/*t* frequencies, averaging 0.32 (SD=0.05) in two-locus models, with natural frequencies ranging from 0.14 to 0.31 in populations in which *t* is not very rare or absent (Ardlie and Silver 1998). While +/*t* frequency was on the high end of naturally occurring frequencies, *α* values above 0.02 led to unrealistically frequent *t* extinctions. Below, we describe the deviations from this natural condition that were also analyzed to understand the effects of both deleterious *t* traits on the outcomes of the simulations.

#### Female multiple mating

To examine how polyandry impacts the divergence between evolving dispersal propensities in + and *t*, we varied *α*, which impacts how likely a female mates with more than one male (see *Mating*), between 0 and 0.02. The number of *t* chromosomes in the simulation is influenced by the local density and *α*, see SI Figure 3 a for this relationship in simulations where all mice disperse with probability 0.1, without any evolution of that trait).

**Figure 3:**
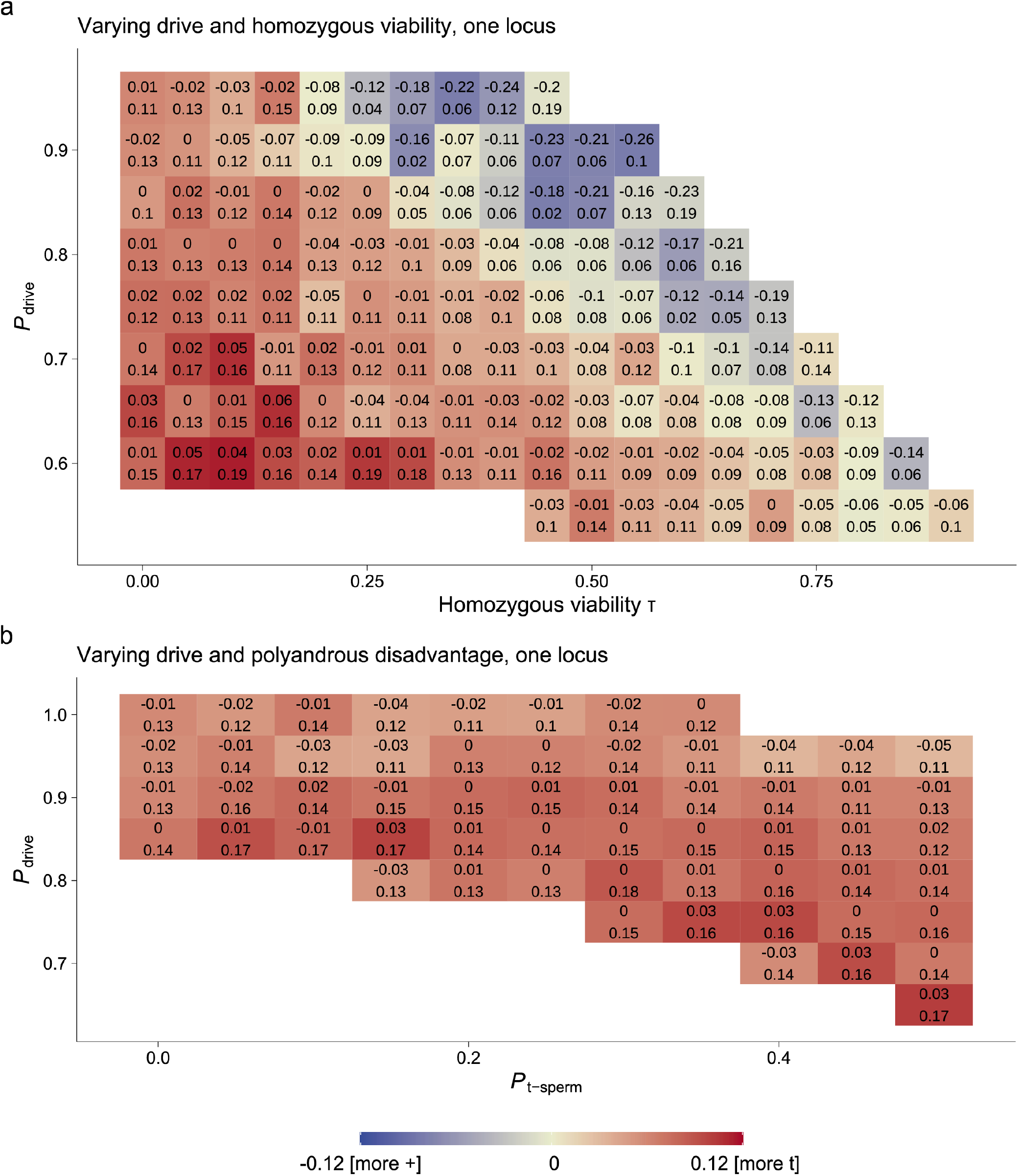
Heatmaps showing dispersal propensity differences between a driver (*t*) and the wild type locus (+) under broadly varying driver traits. Red indicates increased *t* dispersal, blue indicates increased + dispersal. The entire maps of values were tested in 0.05 value increments, white areas indicate that one genotype fixated. The text indicates the 95% confidence interval. **a)** Varying *t* homozygous viability *τ* and *P*_drive_ in simulations without polyandry. **b)** Varying *t* disadvantage in polyandry *P*_t-sperm_ and *P*_drive_ with *α* = 0.02.

#### Homozygous lethality

We also varied *τ*, the probability with which a *t*/*t* embryo is viable, to examine the extent to which *t*’s homozygous lethality is responsible for dispersal evolution. To keep the intended effects operating in our model, *τ* must not be too high as otherwise +/+ will be outcompeted, which eliminates competition between + and *t* which we envisage to be important for the evolution of dispersal in this system. Given that we assume a dominant effect of *t* on dispersal in +/*t*, only +/+ produce the +’s phenotype and their presence is essential for understanding differences between + and *t* in dispersal. We set *τ* to 0.0, 0.25, or 0.5 (for the resulting *t* frequencies for the latter two, see SI Figures 3 b & c).

#### Infertile *t*/*t* males

As described in the introduction, the consequences of *t* haplotype homozygosity can be either inviability or male sterility. While we focused primarily on inviable *t*/*t* (the case for the *t* variant in which dispersal effects were studied in Runge and Lindholm (2018) & Runge and Lindholm (2021)), we also examined conditions in which *t*/*t* were infertile as males, but fully viable (Lewontin (1962); *τ* = 1.0). In this case we assumed that they approach females and mate with them normally, but are completely ignored as potential sires. Some females will, in such a setting, mate with infertile males only, and have no offspring. The resulting *t* frequencies can be seen in SI Figure 3 d.

#### Temporal and spatial heterogeneity

We investigated differences in dispersal propensity under varying environmental heterogeneity (in the otherwise natural condition *α* = 0.02 & *τ* = 0.0) in two-locus simulations. We ran all combinations of spatial heterogeneity 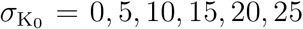 and temporal heterogeneity 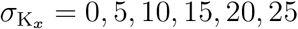 and investigated how these impacted dispersal differences and *t* survival.

#### Weak drive and low fitness costs

To understand if meiotic drivers generally lead to increased dispersal, we searched for combinations of homozygote viability *τ* = 0.5…1 and drive strength *P*_drive_ = 0.5…1 in increments of 0.05 that allowed for long-term survival of both + and *t*, without polyandry, as well as combinations of the proportion of offspring sired by a driver-carrier in a polyandrous mating *P*_t-sperm_ = 0.0…0.5 and *P*_drive_ = 0.5…1, with *τ* = 0; *α* = 0.02. All of these combinations were simulated with 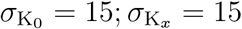. We only did this for one locus models as they provided faster results.

### Execution and analysis of the simulations

We ran simulations for 100,000 (one-locus models) or 1,000,000 (two-locus) turns for each condition (see SI for a table of how many times each condition was run). To visualize the evolved dispersal functions, we combined all simulations with the same conditions, randomly selected up to 100,000 chromosomes per haplotype (*t* or +) in the last 10 turns and extracted each haplotype’s loci. For the one locus models, we then analyzed the distributions of the dispersal propensity *D*_0_. In the two-locus models, we analyzed the dispersal propensities at low density (defined as the mean minus one SD carrying capacity (*±*1), “*µ −* (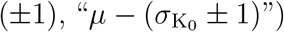 *±* 1)”) and high (*µ* + 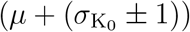) density. These choices allow us to show the variation in dispersal propensity using violin plots.

To quantify the differences in dispersal propensity between *t* and +, we calculated the mean dispersal difference between the genotypes and a 95% confidence interval of this difference using a t distribution with the degrees of freedom equaling the number of individuals that were used as a basis for the difference. Whether and by how much this confidence interval overlapped 0 was used to understand whether the difference in dispersal was clear. Note that all heatmaps use their own color-to-value distribution to ensure best visibility. All presented results are based on fully completed simulations, thereby excluding those where coexistence was not achieved (i.e. *t* or + died out). For an overview of all simulations that were run, including how many did not coexist until the maximum number of turns, see SI Table 1.

## Results

### Dispersal is increased in +/*t* due to its deleterious traits

In one locus models, +/*t* exhibited an increased dispersal propensity compared to +/+ in the natural condition {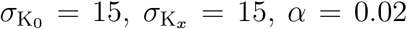, *α* = 0.02, and *τ* = 0}, with an average increase in dispersal propensity of 0.07 (95% CI: −0.01 to 0.15, Figure 2, SI Figure 4). As seen in Figure 2, this difference became smaller with decreasing levels of polyandry (decreasing *α*), and also changed direction as homozygous *t*/*t* became more viable (increasing *τ*) when both deleterious *t* traits were least pronounced (*α* = 0 and *τ* = 0.5). In that condition, +/+ generally dispersed more (−0.07, −0.21 to 0.06).

### The *t* evolves an increased density-dependent dispersal propensity

The two-locus model allows us to investigate differences in the density-dependence of the dispersal phenotype. In the natural condition, +/*t* generally evolved to disperse at much higher rates than +/+ in all densities, but more strongly at high densities (mean difference: 0.14, −0.04 to 0.32; low densities: 0.09, −0.03 to 0.21). This difference was achieved via an increased *D*_1_ (dispersal slope with density), while *D*_0_ (dispersal at 0 density) was essentially identical between *t* and + (Figure 2, SI Figures 5 & 6).

The evolution of this difference again depended on both deleterious traits (Figure 2). However, the results differed from the one locus models. Higher polyandry probabilities *α* increased rather than decreased the difference in dispersal of *t* and +, but at high densities, the confidence interval of the difference in dispersal propensity was consistently smaller with any increase in *α*. Similarly to the one locus models, increases in *t*/*t* viability *τ* changed the direction of the difference in dispersal propensity, with mostly, but not consistently increased confidence intervals. At higher densities, +/+ never clearly evolved to disperse more than +/*t* on average.

When investigating both genotypes separately (SI Figures 7 & 8), *t* was found to evolve the largest high density dispersal when *α >* 0 & *τ* = 0, but once again, a negative relationship between dispersal propensity and *α* can be observed. In contrast, + dispersal propensity from high densities generally increases with increasing *α*, but not above *t*’s propensity.

### Infertile homozygous males also select for increased dispersal

In the case where *t*/*t* males are viable but sterile, the results for the difference in dispersal between *t* and + were overall qualitatively the same (SI Figure 9) as in the case of inviable *t*/*t* described above. However, confidence intervals were larger for sterile *t*/*t* simulations and mean differences were greater (e.g. at *α* = 0.02, the dispersal difference at high densities was 0.19, −0.13 to 0.52 compared to the 0.14, −0.04 to 0.32, mentioned above).

### Spatial heterogeneity is important for *t* survival and dispersal

We also simulated the natural condition in two-locus models under combinations of spatial heterogeneity 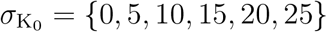 and temporal heterogeneity 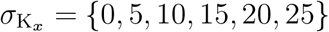. We found that *t* could survive under almost all conditions with the exception of very high spatial heterogeneity 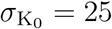 and low temporal heterogeneity 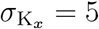. The differences between + and *t* were more pronounced with increasing spatial heterogeneity, but did not have clear pattern with temporal heterogeneity (SI Figure 10).

### Weak drive with some viability costs also promotes dispersal

We found that in 100% (*n* = 27) of all combinations of *τ* and *P*_drive_ that evolved a clear difference between driver and wildtype dispersal (i.e. survival of wildtype and driver, no overlap with 0 in the 95% CI), the driver had an increased dispersal propensity compared to the wildtype (Figure 3 a). This was also the case in 67% (*n* = 98) of simulations without a clear difference. This result was also found in 100% (*n* = 26 for clear differences and *n* = 32 for unclear ones) of the conditions when we varied *t*’s disadvantage in polyandrous matings, *P*_t-sperm_, rather than *τ*, and *P*_drive_ (Figure 3 b). In both cases, visual inspection revealed that the magnitude and direction of the difference was associated with the balance of drive efficacy and costs. The stronger the disadvantage of the driver, the more it increased dispersal.

## Discussion

Our results show that the *t* haplotype evolves a more dispersive phenotype than the wildtype under conditions that mimic natural settings (i.e. with polyandry and full *t* homozygous lethality), particularly at high densities. By comparing the natural condition of the *t* haplotype with hypothetical conditions where we varied the female mating rates and the viability and fertility of *t*/*t*, we were able to demonstrate that all of *t*’s known disadvantages are jointly responsible for the elevation in dispersal propensity. However, higher multiple mating frequency did not consistently result in larger dispersal differences. Moreover, by varying driver efficiency, homozygous costs, and polyandrous disadvantages, we showed that driver disadvantages more generally select for increased dispersal.

All of the modeled *t* disadvantages (polyandry, homozygous lethality, and homozygous male infertility) contributed to increased dispersal of +/*t* or *t*/*t* over +/+, both at high and low densities. At full homozygous lethality, no polyandry was needed to produce a clear increase of *t* over + dispersal in one or two locus models. In contrast, at 25% homozygous viability, only models with polyandry showed a conclusive increase in dispersal in one locus models, but no polyandry was needed to show a dispersal difference when density-dependent dispersal was possible. Thus, when *t* can evolve a density-dependent dispersal propensity, the dispersal rates diverged from + already at smaller *t* disadvantages. This is likely in part a result of our assumption that polyandry increased with density, a pattern also found in studies in the field (Dean et al. 2006; Firman and Simmons 2008; Manser et al. 2020). However, polyandry did not increase the differences in dispersal in high densities with low *t*/*t* viability; under such settings, +/+ individuals, too, increased their high-density dispersal with increasing polyandry. Clearly, +/+ also benefit from avoiding polyandrous matings and thus engaging in sperm competition, likely with their kin, but to a lesser degree than *t*-carriers. Nonetheless, +/+ were only as dispersive or more dispersive than *t*-carriers when at least one negative *t* trait was reduced.

We also found that *t* and + could co-exist in a wide range of spatially and temporally heterogeneous (or homogenous) environments. Yet, moderate spatial heterogeneity was required to elicit a strong differentiation between +/+ and +/*t* dispersal at high densities. No clear impact on dispersal differences could be found for temporal heterogeneity. In nature, house mice are widespread and live in very heterogeneous habitats, from less dense and more temporally heterogeneous feral to dense, more stable commensal populations (Bronson 1979). Thus, natural habitats of house mice are spatially and temporally heterogeneous and likely fulfill the requirements for the evolution of dispersal differences.

In the alternative setting where *t*/*t* were fully viable, but male *t*/*t* were infertile, *t* frequencies evolved to be much higher, but the difference in dispersal between *t* and + evolved to the same qualitative pattern as in the *t*/*t* inviability scenario. Quantitatively, the difference in dispersal was higher with infertile *t*/*t* males, especially at high densities, but less convincing due to larger confidence intervals. Both scenarios imply that matings between *t*-carriers are deleterious and should be avoided, and dispersal helps to achieve this. Similarly, in both cases, remaining philopatric carries an increased risk of local extinction: for inviable *t*/*t* this could happen if genetic drift is strong, as *t* frequency is limited to 0.5 and can only drift to 0, not 1; and for infertile *t*/*t* as their fixation would crash their population. Infertile *t*/*t* males compete for resources with +/*t* males, which could lower *t* fitness, which has been speculated to select for inviable *t*/*t* (Silver 1993). However, we found that *t* frequency was much higher when *t*/*t* were infertile, but viable, with either polyandry or dispersal evolution keeping *t* frequency at bay. An increased dispersal propensity could alleviate the negative fitness consequences of male infertility by avoiding deleterious matings, which may reduce selection towards inviability, making the latter more likely a by-product of reduced recombination (Sugimoto 2014) rather than a selected trait when dispersal is high enough.

Finally, we also modeled driver dispersal more generally by investigating evolved dispersal differences between driver-carriers and wildtypes for two types of drivers: drivers with varying viability and drive strength in populations without any polyandry, and drivers with homozygous inviability, but varying drive strength and disadvantage in polyandrous matings. Whenever coexistence of driver and wildtype was possible, i.e. the driver was neither too weak nor too successful, drivers were generally selected to increase dispersal. Essentially, the less successful the driver, the more it was selected to disperse. Drivers that are not fixated or close to fixation should roughly fit into this category of not being ‘too successful’ (Price et al. 2019). The wildtype was also selected to disperse when the driver was close to being too successful for coexistence, but in these conditions confidence intervals of the dispersal difference were always overlapping 0, thus our results did not predict a clear difference in dispersal. Based on these results, we predict that meiotic drivers that are genetically linked with increased dispersal should outcompete drivers that are not. Thus, it would be interesting to test other systems for the presence of increased dispersal of driver carriers, which has so far not been done.

Previous models of the *t* haplotype’s dispersal phenotype, derived during a time when empirical evidence was unavailable, predicted that dispersal was particularly important for *t*’s fitness, either because wildtype-fixed populations should be easily infected by *t* (Lewontin and Dunn 1960), or because sub-populations carrying the *t* would go extinct frequently (Lewontin 1962), or because a reduction in +/*t* dispersers due to selection between sub-populations would lead to low *t* frequencies (Nunney and Baker 1993). These studies, however, could not consider the *t*’s disadvantage in polyandrous contexts because it was only discovered later (Olds-Clarke and Peitz 1985; Manser et al. 2011; Sutter and Lindholm 2015). Ours is the first model that includes the effects of both deleterious traits on the dispersal phenotype. Our results indicate that including these traits is essential for understanding the evolution of dispersal differences between the *t* haplotype and the wildtype.

We did not include potentially sex-specific dispersal phenotypes in our model; for example, one could speculate that only +/*t* males should disperse at higher rates because of the problems their sperm encounter in multiple mating contexts. We chose this simplifying assumption primarily because we did not see sex biased effects in our long-term field study (Runge and Lindholm 2018). Our current results show that a potentially more easily evolvable, sex-independent effect can evolve. It is also conceivable that females, as mothers of some +/*t* sons, may profit from moving to places where the *t* haplotype tends to do well. To that end, a study that asked very different questions from ours has found that fitness benefits of dispersal that are reaped a few generations after a dispersing ancestor can still select for dispersal (Travis et al. 2009). Either way, selection towards increased dispersal of +/*t* females is likely weaker than on +/*t* males. Our results, with clear differences in dispersal phenotype between +/+ and +/*t* when dispersal effects were constrained to be identical for both sexes, reflect the result of selection that is averaged over the two sexes.

There is an ongoing effort to create a male-determining-gene-carrying *t* haplotype drive system (*t*-*SRY*) to eradicate harmful house mouse populations (Piaggio et al. 2017; Kanavy and Serr 2017; Gemmell and Tompkins 2017). It is crucial for the safety and success of this project to understand the dynamics of the *t* in the wild (Manser et al. 2019). In this study, we have provided evidence that *t*-carrying mice can be expected to have an increased dispersal propensity, which could help them spread a modified *t* haplotype further than planned. It is therefore important to model the influence of increased dispersal when considering the impact of the *t*-*SRY* system in the wild.

Our study provides, to our knowledge, the novel result and explanation of how an intragenomic conflict involving a meiotic driver can select for increased dispersal of driver-carrying individuals. Changes in behavior of driver-carriers have so far rarely been documented. A comparable phenomenon is found in fire ants where colonies of ants carrying a driving supergene are differently organized than those of non-carriers (Wang et al. 2013; Ross and Shoemaker 2018)). In general, parasites (that are not drives) often benefit from increasing the dispersal rate of their hosts (Lion et al. 2006) or increasing their mating rate (e.g., the increased mating rate of *Wolbachia*-infected flies (Champion de Crespigny et al. 2006)). In summary, we showed how drivers can evolve an increased dispersal of their carriers. With this, we add another layer to the already complex intragenomic conflict between the driver and the rest of the genome.

## Supporting information

Data frame showing total counts of simulations run for each condition

## Acknowledgements

We thank Andrés Bendesky and Barbara König for computing resources during parts of this study and Andri Manser for feedback on an earlier version of the manuscript.

## Author contributions

JNR conceived the study, programmed the simulation, and analyzed the data. JNR and HK contributed to simulation design. JNR, HK, and AKL wrote the manuscript.

## Data availability

The code of the simulation is available at https://github.com/jnrunge/t-vs-w-dispersal-evolution/. The data will be available as soon as it is compressed and uploaded at https://doi.org/10.5281/zenodo.4486286.

