## Supplemental figures for "Selfish migrants: How a meiotic driver is selected to increase dispersal"

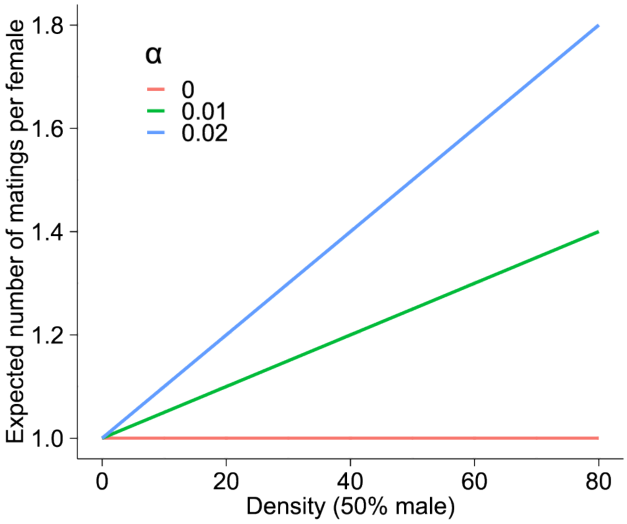


**Suppl. Fig. 1:** Expected number of matings per female with increasing density at different $\alpha$ values.

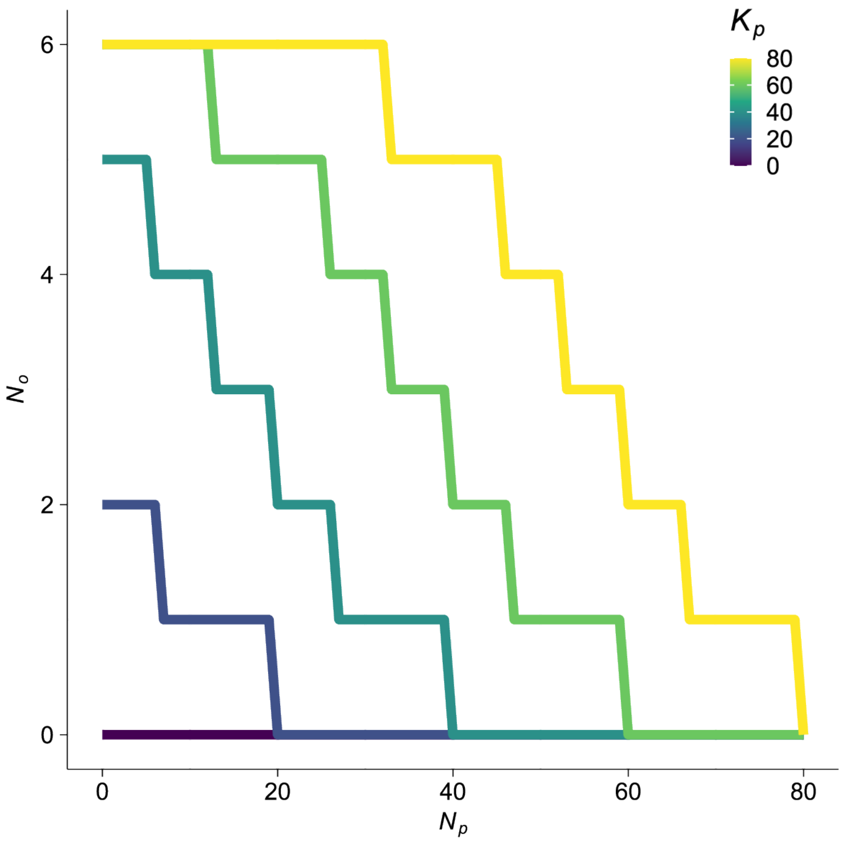


**Suppl. Fig. 2:** Female fecundity at different densities and carrying capacities.

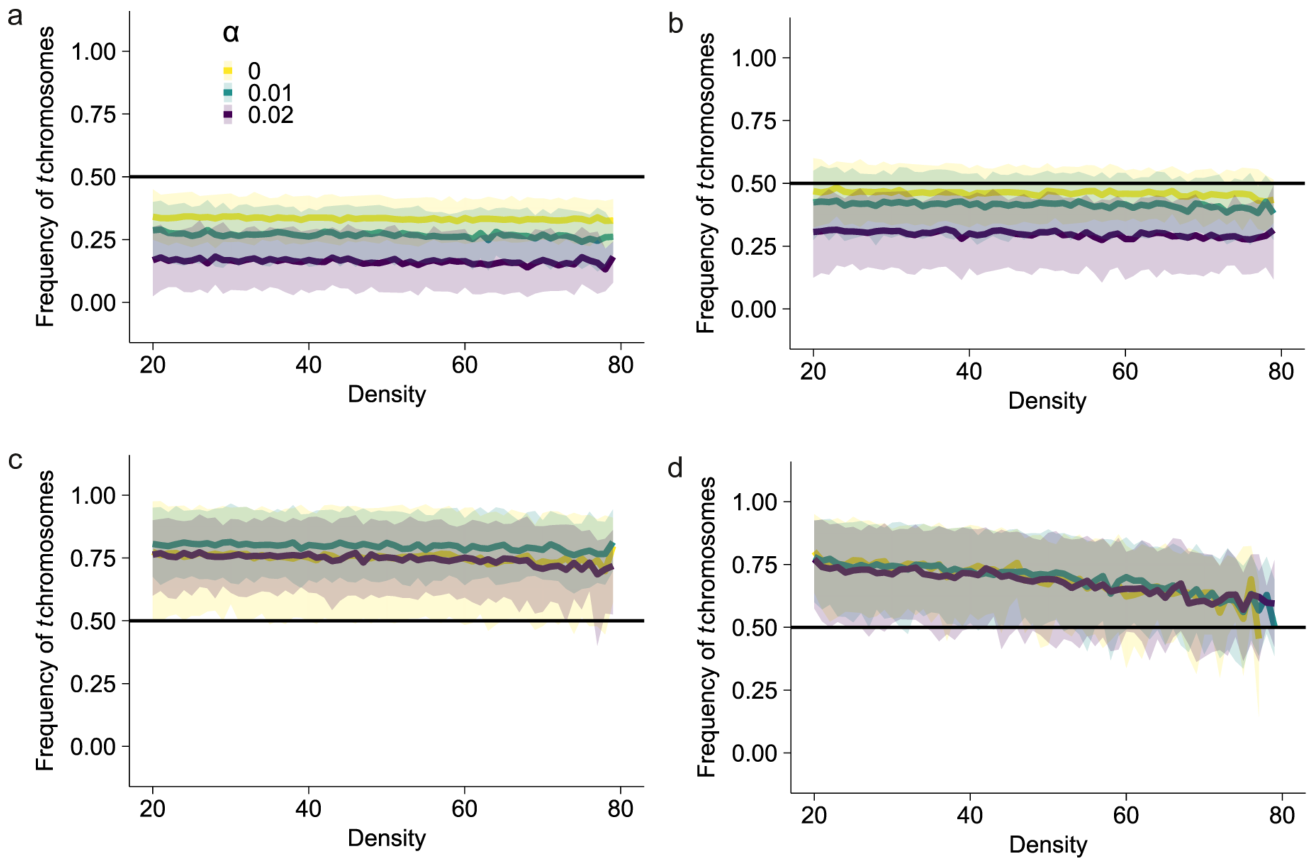


**Suppl. Fig. 3:** Frequency of $t$ chromosomes (0.5 in $+/t$, 1 in $t/t$) in simulations after 10,000 turns of $t$ and $+$ coexistence with a fixed dispersal propensity of 0.1 and varying $\alpha$ under $\tau = 0.0$ (a), $\tau = 0.25$ (b), $\tau = 0.50$ (c), or infertile male $t/t$ (d, $\tau = 1$). Lines indicate the mean and ribbons show the standard deviation.

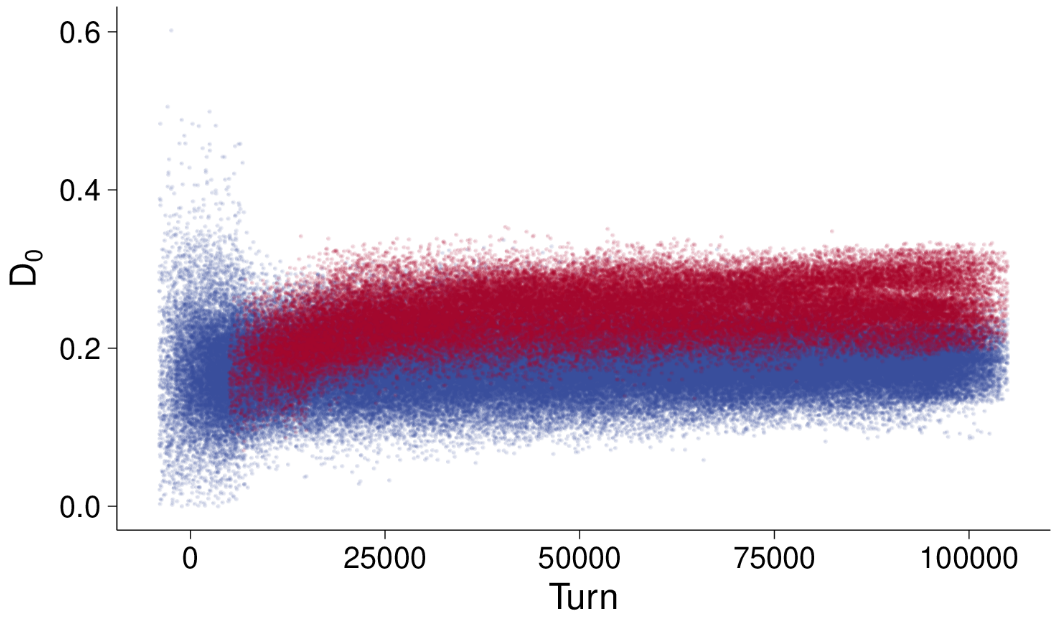


**Suppl. Fig. 4:** Evolution of dispersal propensity $D_0$ over the course of the simulation under natural conditions with one locus. A random subset of individual values is visible as dots, jittered horizontally. $t$ values are in red, $+$ values are in blue.

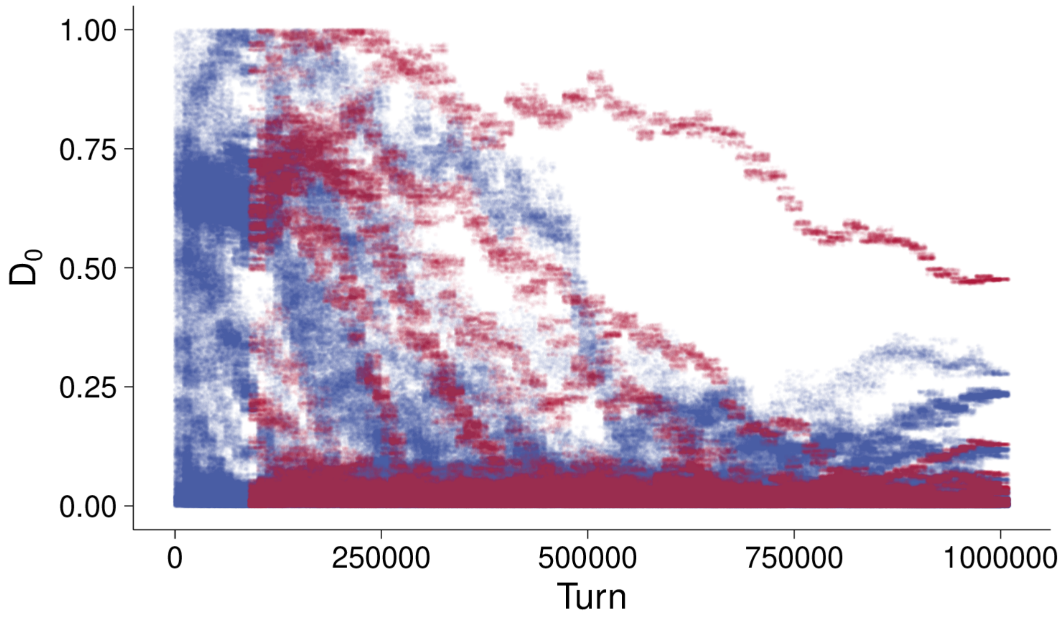


**Suppl. Fig. 5:** Evolution of dispersal propensity intercept $D_0$ over the course of the simulation under natural conditions (two-locus models). A random subset of individual values is visible as dots, jittered horizontally. $t$ values are in red, $+$ values are in blue.

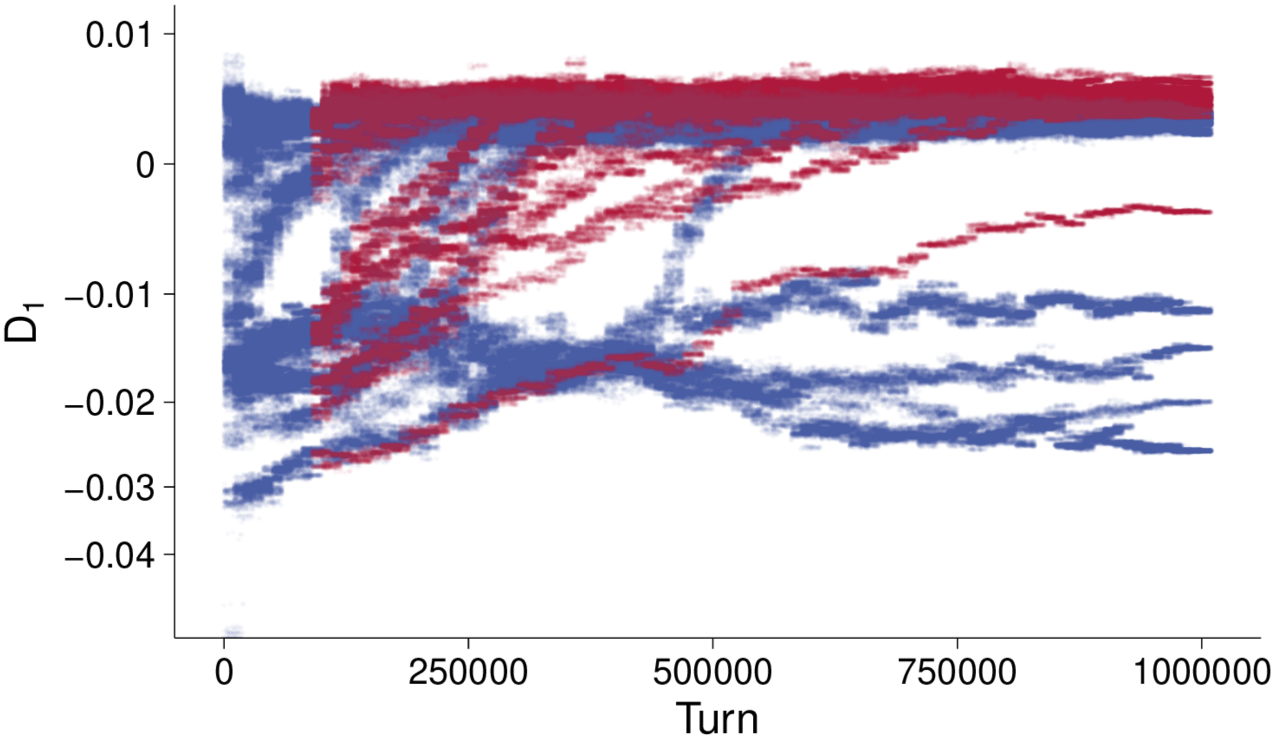


**Suppl. Fig. 6:** Evolution of dispersal propensity slope $D_1$ over the course of the simulation under natural conditions (two-locus models). A random subset of individual values is visible as dots, jittered horizontally. $t$ values are in red, + values are in blue.

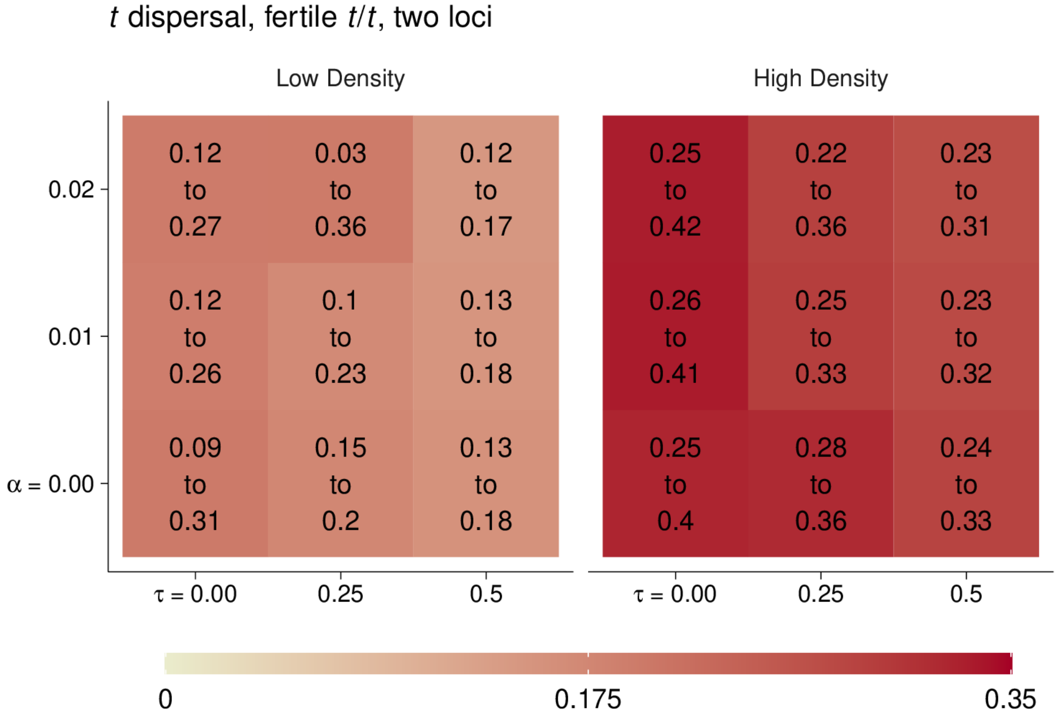


**Suppl. Fig. 7:** Heatmap showing the mean dispersal propensity of $t$ in two-locus models in low and high densities in varying polyandry $\alpha$ and $t/t$ viability $\tau$. Red indicates increased $t$ dispersal. The text indicates the 95% confidence interval.

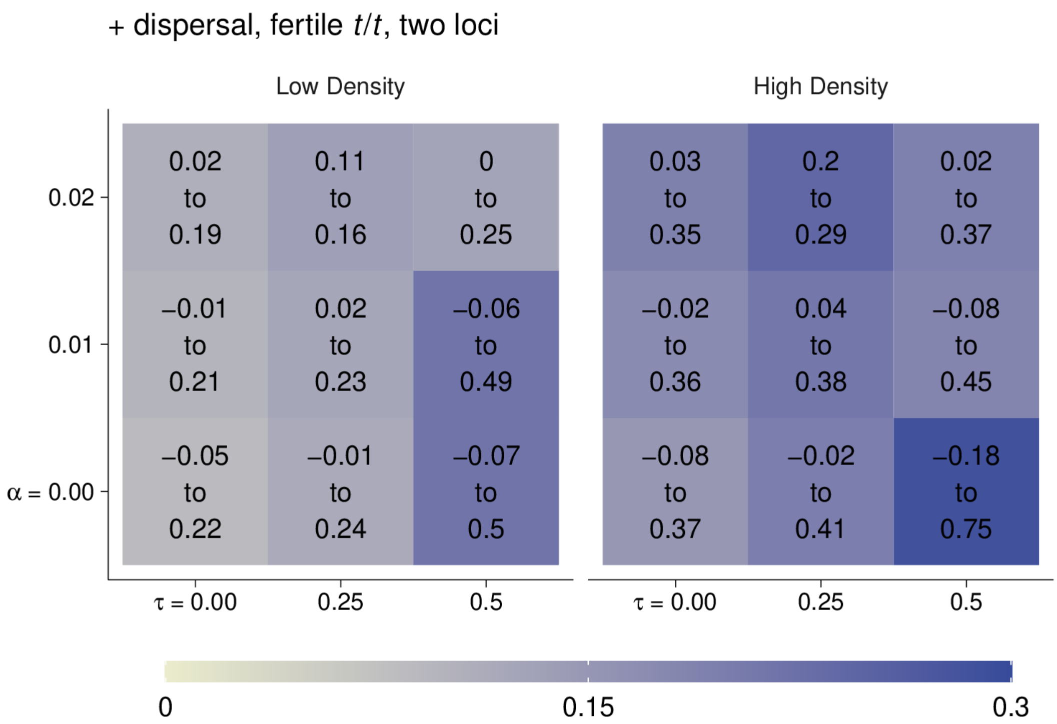


**Suppl. Fig. 8:** Heatmap showing the mean dispersal propensity of + in two-locus models in low and high densities in varying polyandry $\alpha$ and $t/t$ viability $\tau$. Blue indicates increased + dispersal. The text indicates the 95% confidence interval.

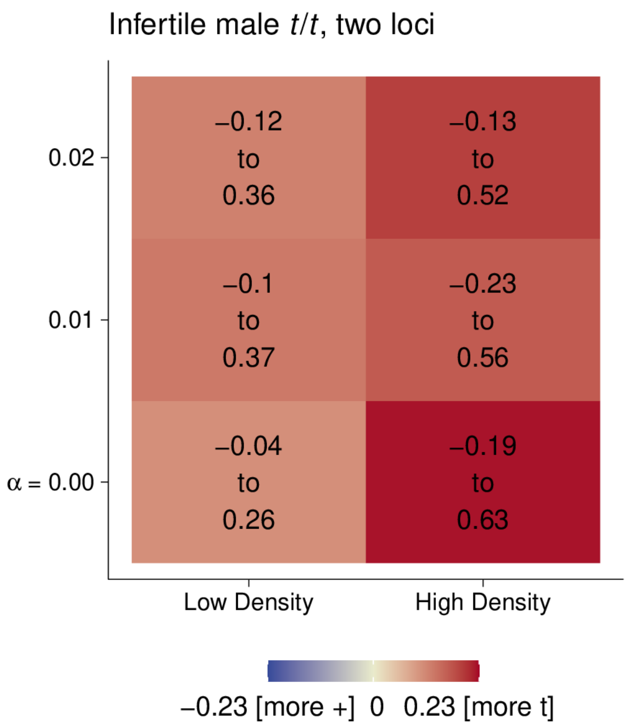


**Suppl. Fig. 9:** Heatmap showing the mean difference between $t$ and + dispersal propensities in two-locus infertile male $t/t$ models in low and high densities under varying polyandry $\alpha$. Red indicates increased $t$ dispersal, blue indicates increased + dispersal. The text indicates the 95% confidence interval.

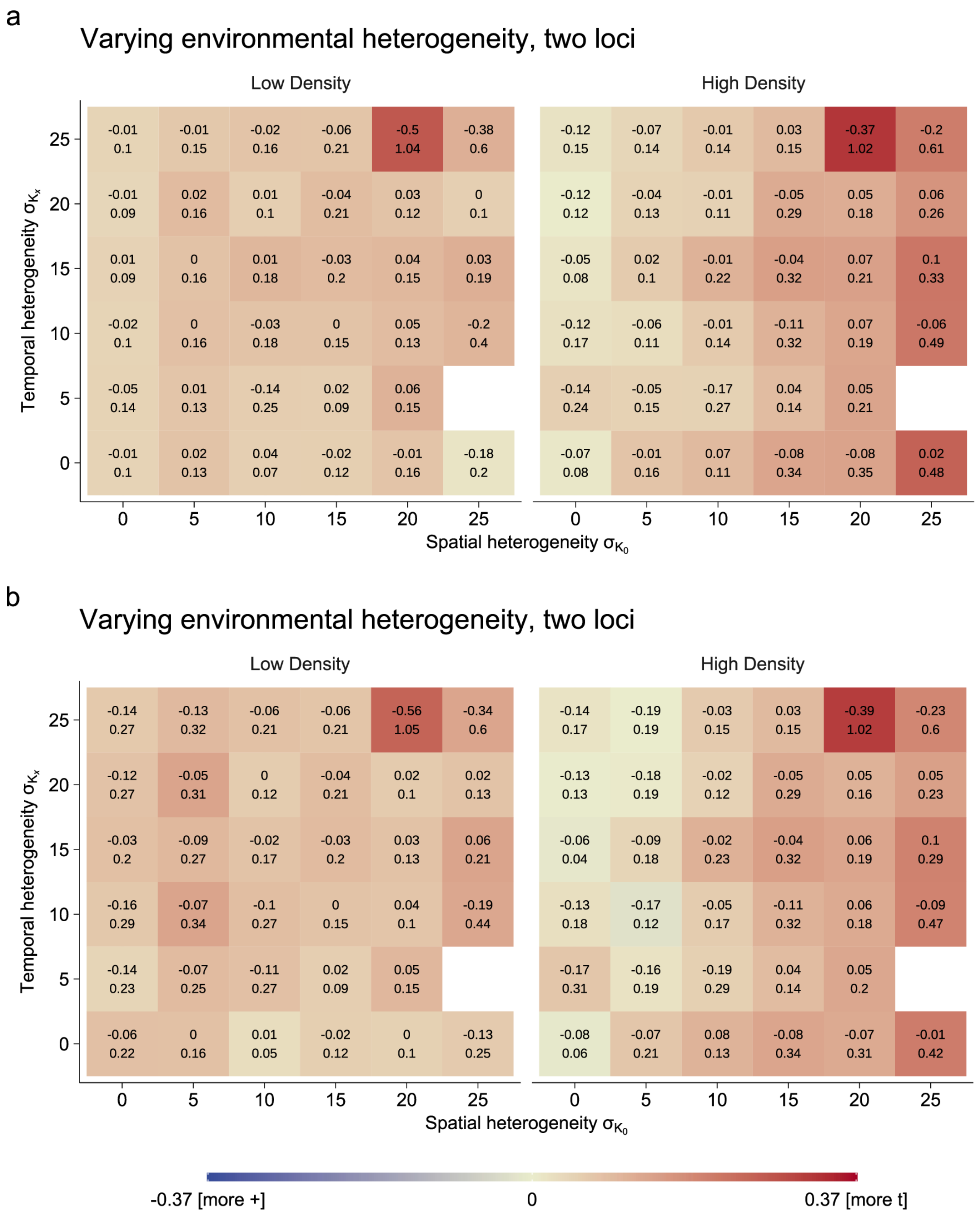


**Suppl. Fig. 10:** Heatmaps showing the mean difference between $t$ and + dispersal propensities in two-locus natural condition models in low and high densities under varying spatial and temporal carrying capacity heterogeneity. Red indicates increased $t$ dispersal, blue indicates increased + dispersal. White areas indicate that + fixated, the text indicates the 95% confidence interval. **a)** What is considered high/low density is calculated based on the individual (x axis) $\sigma_{K_0}$ levels. **b)** High/low density is based on $\sigma_{K_0} = 15$ like in the main text.
